# The effect of wild *Saccharomyces* on composition and aroma of the Sauvignon blanc

**DOI:** 10.1101/2020.08.03.214437

**Authors:** Sandra D. C. Mendes, Stefany Grützmann Arcari, Mauricio Ramirez-Castrillon, Simone Silmara Werner, Patricia Valente

## Abstract

*Saccharomyces* strains isolated from vineyards in Southern Brazil were used as starter in micro-fermentations of Sauvignon Blanc juice (SB) to study the ability to produce different aromatic profiles. The molecular characterization allowed to differentiate the isolates from vineyards 06 CE, 11CE, 33CE, 01PP, 12M, 13PP, 26PP, 28AD, 41PP from the reference strain (2048SC) used in wine production. Under the same conditions tested, each strain belonging to the genus *Saccharomyces* produced metabolites in different concentration and combination, significantly influencing the aroma of the SB must. Volatile compounds such as octanoic acid, diethyl succinate and ethyl lactate were associated with the strains 26PP, 41PP, 01PP and 12M, while the strains 33CE, 28AD, 13PP and 06 CE were associated with the production of ethyl acetate, 1-hexanol, highlighting 06CE for production of 1-hexanol (592.87 ± 17.46 µg/L). In addition, the olfactory activity value (OAV> 1) allowed the evaluation of the range of participation of each compound in the final SB fermented. Finally, the selected wild strains are promising to improve the regional character of the wine.

## INTRODUCTION

Human civilizations continually look for to control the fermentation process of beverages and food by progressively selecting specific yeasts adapted to their needs. The use and reuse of yeast over the centuries resulted in the emergence of these strains due to the selective pressure of the process (1). In the case of domestication, it is expected that the differentiation of certain phenotypic traits is accompanied by genetic differentiation (2). Thus, the reduced levels of variation present in the strains used for wine fermentation was the result of a genetic bottleneck by selection of specific characteristics. These characteristics include the fermentation of sugars present in the must (<2.0 g/L residual sugar), low H2S production and tolerance to osmotic and ethanol stress, tolerance to temperature variation and SO2, low foam production (3, 4). With the advent of metagenomics studies, industrial yeasts represent a small fraction of the universe of existing yeasts, (5, 6). Thus, it became interesting to isolate new yeast species or different *Saccharomyces cerevisiae* genotypes that generate different flavor profiles in the final product. In this context, ‘Sauvignon blanc’ juice was used as a model system and the strains of the genus *Saccharomyces* selected from vineyards from regions of Santa Catarina as a starter to demonstrate the performance of these strains in the phenotype of the final product, wine. A complex mixture of chemicals was studied, mainly ethyl esters, higher alcohols, fatty acids both of great relevance in terms of quality and impact for the typicity and denomination of origin in this traditional region.

## RESULTS

### Selection and identification

The isolates were collected from different points of the vineyards and cultured in 2 % YPD medium for 72 hours at 30 °C. Then, the isolates were identified by sequencing D1 / D2 domain of the major ribosomal Subunit (LSU), or ITS1-5.8S-ITS2 region (7). We used species-specific primers ScerF2 and ScerR2 to identify and distinguish *S*. *cerevisiae* species from other species belonging to the genus *Saccharomyces*, which includes S. *bayanus, S*. *pastorianus, S kudriavzevii* found in fermented musts ((8)Muir et al., 2011). Of the nine isolates belonging to the genus *Saccharomyces*, only six (6) were identified by species-specific primers as *S*. *cerevisiae*, with PCR products length of 150 bp. PCR assays using the Intron Splice Site primer EI-1 were used with these strains, three *Saccharomyces* spp (06 CE, 11CE, and 33CE) showed a different profile from the rest of the vineyard isolates. Thus, PCR fingerprinting technique using the EI-1 primer was able to group members of the species *S*. *cerevisiae* but were not suitable to resolve the hybrid isolates. Strains 06CE, 11CE, and 33CE did not assimilate α, α-trehalose, maltose or raffinose differing from the description of the species but cannot be considered a potential new species.

### Determination of volatile compounds before fermentation

The volatile compounds that constitute the aroma of wine are traditionally divided into three classes according to source: grape aroma, fermentation, and maturation. A total of 44 volatile compounds were identified in the SB must from an altitude vineyard in the region of São Joaquim, South Brazil (Table S1). Of these, 20 volatile compounds were positively identified using commercial standards. The other compounds were tentatively identified based on the similarity between the mass spectrum of the sample compounds and the NIST library (similarity> 70%), as well as, through the calculated retention index, against the retention index found in the literature, which considered the values reported for polar columns of polyethylene glycol. A maximum deviation of 36 units was observed between the experimental data and the retention index reported in the literature for the benzaldehyde compound and from 29 units for 3-hexen-2-one and isocitronelol. From the chemical groups found in the volatile fraction of the SB juice, the esters were present in a larger number (12 compounds), followed by the higher alcohols (11), acids (5), ketones (4), C13-norisoprenoids (4), aldehydes 3), terpenes (3), phenol (1), and lactone (1).

### Determination of volatile compounds after fermentation

Grape-derived compounds such as terpenes, pyrazines and thiols are known to play a key role in the aroma of SB, while the alcoholic fermentation conducted by *S*. *cerevisiae* induces the formation of active secondary aroma metabolites such as ethyl esters, higher alcohols and fatty acids, as demonstrated by Principal Component Analysis. The two first principal components obtained considering the chemical groups in volatile fraction explained 71.50 % of the total variance of the data (Fig1). The chemical groups such as terpene, fatty acid and ethyl ester had a positive effect on the first PC, while the ethyl ester and higher alcohol groups had positive effects on the second PC. The strain 11CE and the SB juice have significant negative factor loadings in component 1and 2 (Fig.1a and Table S2, Table S3). The second component explained 23.73% of the data variance and was formed by lactone group. Figure 1a shows how 26PP, 12M, 41PP and 01PP have significant positive factor loadings in component 1, and are associated with the production of terpene, fatty acid and ethyl ester; while 06CE and 13PP have significant positive factor loadings in component 2, associate with the production of higher alcohol and acetate ester, highlighting the strain 06CE (Fig. 2a) for the production of 1-hexanol (592.87 ± 12.35 μg/L) and the production of ethyl acetate (7574.84 ± 1786.28 μg/L), furfuryl acetate (25.54 ± 4.93 μg/L) and S-furfuryl thioacetate(1.80 ± 0.0 μg/L) (Fig. 2b). The major production of β-damascenone (27.64 ± 0.61 μg/L) and β-ionone (1.71 ± 0.0 μg/L) was observed in the strain 01PP (Fig. 2c).

**Figure 1.**
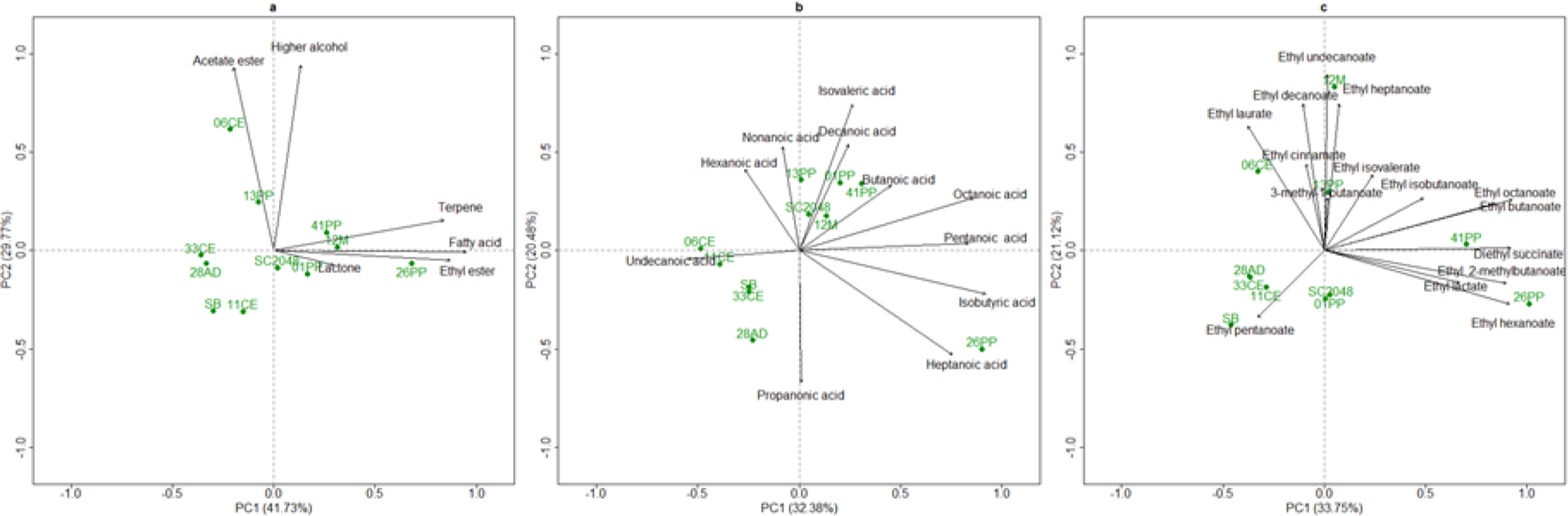
Principal components analysis of volatile compounds in yeasts **a**. Compounds produced during the micro-fermentation of Sauvignon Blanc (SB). Inoculation with a commercial strain of *S*. *cerevisiae* (2048SC) and wild strains (26PP, 41PP, 01PP, 12M, 33CE, 28AD, 13PP, 06 EC and 11CE), including must (SB) **b**. Principal components analysis of fatty acids **c**. Principal components analysis of ethyl esters

**Figure 2.**
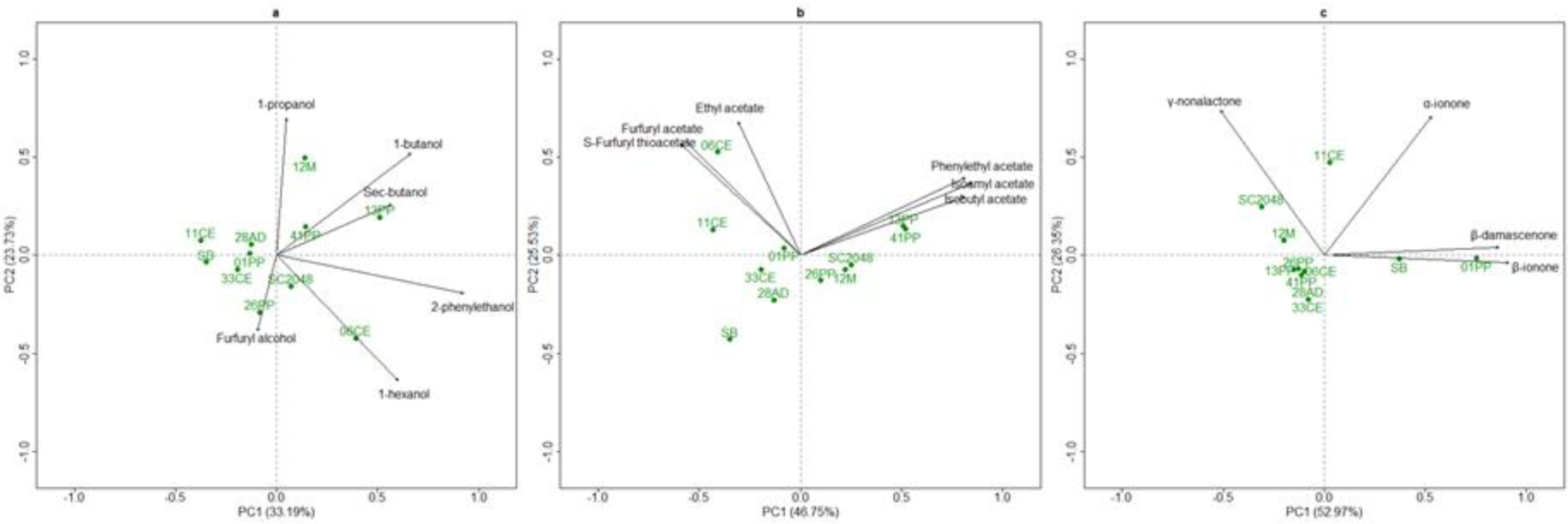
Principal components analysis of volatile compounds in yeasts **a**. Compounds produced during the micro-fermentation of Sauvignon Blanc (SB). Inoculation with a commercial strain of *S*. *cerevisiae* (2048SC) and wild strains (26PP, 41PP, 01PP, 12M, 33CE, 28AD, 13PP, 06 EC and 11CE), including juice (SB) **a**. Principal components analysis of higher alcohols **b**. Principal components analysis of acetate ester c. Principal components analysis of lactone

Fig. 3 shows the compounds perceptible by the human nose and were grouped into three clusters. Furthermore, the system indicates the relative levels of volatile compounds produced by each strain and the importance of the compounds to differentiate their profile from commercial strain SC2048. Those that stand out among higher alcohols are 1-hexanol, 2-phenylethanol, 1-butanol, and among the esters of fatty acids are the compounds ethyl-2-methyl butanoate, ethyl butanoate, ethyl cinnamate, ethyl heptanoate, ethyl isobutanoate, and ethyl isovalerate. In the case of the acetate esters, only phenylethyl acetate showed the concentration that exceeded OAV> 1. The α-ionone, β-damascenone and γ-nonalactone compounds belonging to the class of C13-norisoprenoids also stood out; these compounds, even at low concentrations, contributed positively to the SB flavor.

**Figure 3.**
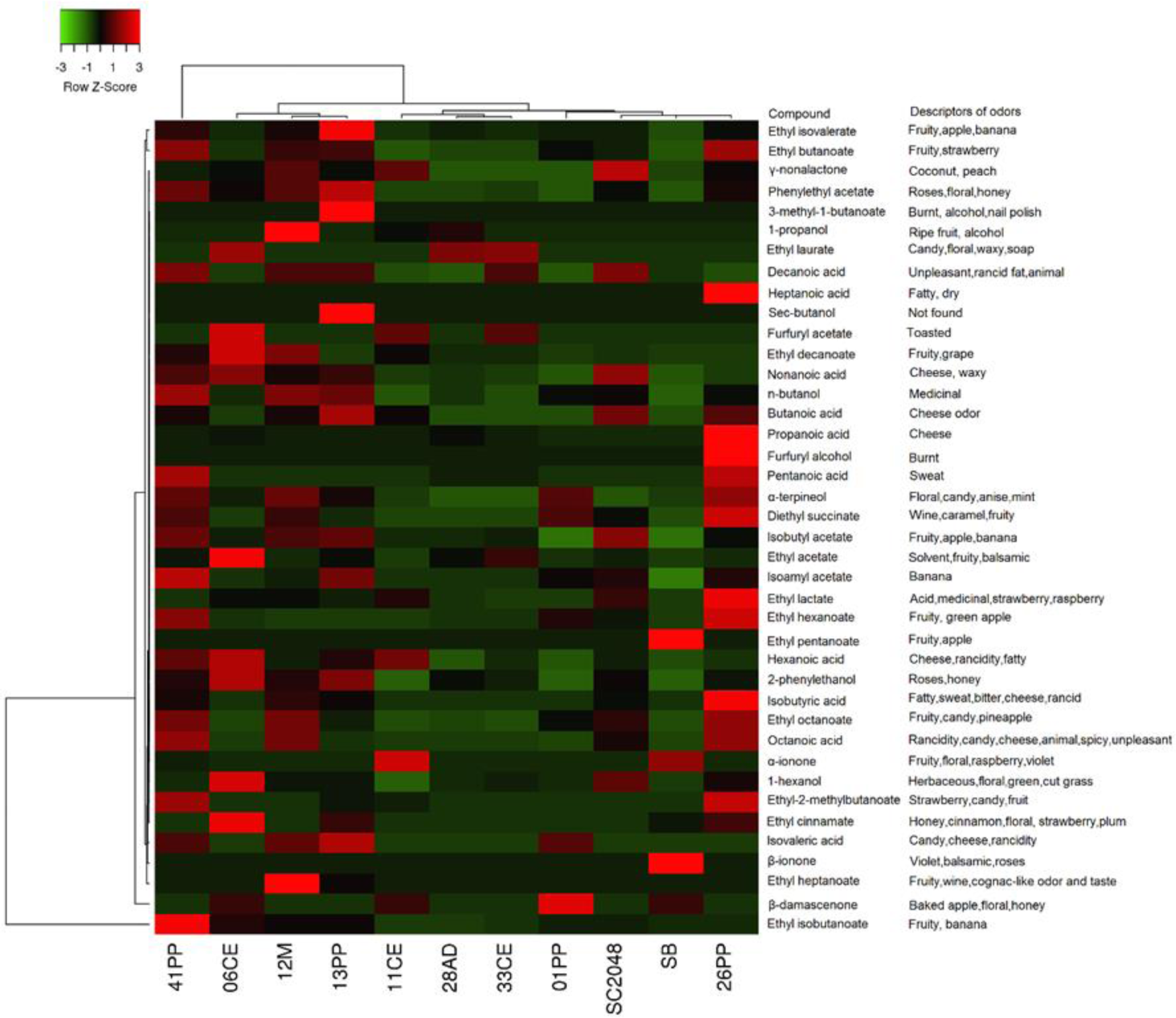
Heat map was generated based on the z-scores representing the level of olfactory activity value of the volatiles produced by wild strains 26PP, 41PP, 01PP, 12M, 33CE, 28AD, 13PP, 06 EC and 11CE and controls (SB must and commercial strain 2048SC). The values are represented in red, high olfactory activity, and green, low olfactory activity, black without a change in the aroma perception of the volatile compounds produced and their descriptors of odors

## Discussion

The identification methods are, in most cases, the first to be applied to reduce the number of interesting strains to be evaluated. rDNA regions (D1 / D2 domain and ITS region) are not useful in the identification of species within the *Saccharomyces* genus, because the sequences do not differ significantly. In the multiplex test using specific primers (ScerF2 and SceR2) to identify nine strains, previously identified as *Saccharomyces* by sequencing the LSU and/or ITS regions, only three strains were not identified as *S*. *cerevisiae*. According to Šuranská et al. (9) and Muir et al. (8), the isolates may be hybrids of *S*. *cerevisiae* and other species belonging to the genus *Saccharomyces*. *Saccharomyces* clade can be found in natural environments and in industrial fermentations as interspecific hybrids between *S*. *cerevisiae, S*. *kudriavzevii, S*. *uvarum* and *S*. *eubayanus* (10, 11). These differences include copy number and ploidy variations, genome rearrangements, and polymorphism changes (12). In the next step of identification, the Intron Splice Site primer EI-1 was proposed to detect polymorphisms between the commercial yeast (2048SC) and strains 33CE, 11CE, and 06CE isolated from vineyards, which were not identified as commercial yeasts *S*. *cerevisiae*. The observed variable traits can be exploited as non-molecular markers in intraspecific diversity studies. However, the presence of industrial strains in the microbiota leads to the reduction of the genetic differentiation between populations. The differences in observed diversity between mtDNA-RFLP and microsatellite analysis may result from the smaller number of bands detected in the profile.

There is a strong influence of the composition of the grape on the formation of compounds of active aroma by the species belonging to the genus *Saccharomyces*. In many cases, the role of yeast metabolism is the release of an aromatic compound from a non-volatile precursor molecule. The genetic profile of *S*. *cerevisiae* is relevant in the formation of metabolites that confer specific aromas of wine. However, several other factors also affect the spectrum of compounds formed(13, 14). Because of this, we characterized the SB juice as demonstrated in (Table S1), before being submitted to the micro-fermentations with the selected strains. We observed the C6 alcohols 1-hexanol, 2-hexanol, (E) -3-hexen-1-ol and (Z) -3-hexen-1-ol, present in the must/juice due to the action of enzymes on linoleic and linolenic acids, extracted from the grapes during the crushing stage. The 1-hexanol molecule is one of the most representative compounds for the aroma of Sauvignon Blanc wines from Victoria, Australia, and Marlborough, New Zealand (15, 16). Changes in volatile wine composition may occur as the yeast responds to the change in the composition of the compound or may reflect changes in the concentration of volatile wine precursors derived from grape (17). The accumulation of compounds is related to the anabolism of grape precursors, which stimulate the production of volatile compounds derived from yeast (18). These compounds may imply the presence of medium chain fatty acids (MCFAs) derived from the grape, or their precursors, contributing to the group of ethyl esters released from MFCA during fermentation, which could provide an important link between the composition of the grape and the volatile wine profile, a product of lipid metabolism(19–21). Fig 1a and b additionally show the vectors indicating the direction and magnitude of the influence that each strain has on the positioning of the compounds produced. Hyma et al. (2) have demonstrated that the divergence in aroma and flavor of the wine is combined with the genetic divergence between wine and wild yeast. Fig. 3 allowed evaluating the scope of participation of each compound at the final flavor, with some substances providing pleasant notes, while others may have negative contributions (22). However, the aroma of the wine is influenced by complex interactions between several components and is seldom dominated by a single component (23, 24). These observations are consistent with (6), who detected a significant effect of selected strains on wine phenotypes in each region.

## Conclusion

In the study, mainly 41PP and 26PP lineages were very promising and demonstrate that the choice of inoculated, commercial yeast or the selected strains belonging to the *Saccharomyces* genus affect the composition and sensory properties. In addition, there are marked differences in perception as presented by the value of olfactory activity (OAV).

## MATERIALS AND METHODS

### Strains used, characterization and extraction of genomic DNA

The strains *Saccharomyce*s (n=8) used in this study were isolated from leaves, bunches of grapes, soil, Strain enrichment and isolation was done as previously described and a commercial strain (*Saccharomyces cerevisiae* CLIB 2048). All strains were stocked at -30 °C in 20 % glycerol and grown in complete YPD broth (0.5 % yeast extract, 2 % peptone, 2 % dextrose). The identification was performed by comparing sequences from two regions: D1 / D2 domain of the major ribosomal subunit (LSU), or ITS1-5.8S-ITS2 region (25). Also, a species-specific primer set was used for confirmation of species of the genus *Saccharomyces*, described by Muir et al. (8). Discrimination was performed using the Intron Splice Site primer EI-1 (5’-CTGGCTTGGTGTATGT-3) (26).

### Determination of volatile organic compounds

#### Preparation of strains belonging to the genus *Saccharomyces*

Sauvignon blanc (SB) must from a highland vineyard in the São Joaquim region, South Brazil (28°16’30”S, 49°56’09”W, alt 1.400 m), was previously treated with 29.0 g/L of sulfur dioxide as sodium metabisulphite (Na2S2O5), 8.0 mL/L of the pectolytic enzyme for clarification of the must, and 7.0 mL/L of bentonite to facilitate the sedimentation of non-fermentable solids, and was transferred to the cold storage chamber for 7 days. After clarification, 200 mL of SB must was sterilized with nitrocellulose membrane (0.22 µM, 47 mm in diameter). A volume of 4.9 ml of SB must was aseptically transferred to vials of 20 mL. The inoculum of each strain was previously cultured at 28 °C for 24 hours in YPD broth. The exponentially growing yeast cells were used to adjust the final cell number to 2.0 ×107 cells/mL. Thereafter, 100 μL of each strain was inoculated into each vial containing 4.9 mL of SB must then incubated at 28 °C for 48 hours.

### SPME fiber preparation

SPME fiber made of DVB/CAR/PDMS obtained from Supelco (Bellefonte, PA, USA) was initially conditioned according to the manufacturer’s recommendations. To each vial containing the fermented SB 1.5 g NaCl was added. The SB fermented/wine was incubated for 5 minutes at 56 °C and then the fiber was exposed in the headspace (HS) for 55 minutes. The adsorption on the injector of the gas chromatograph was performed for 2 minutes at a temperature of 265 °C in splitless mode.

### Extraction of volatile organic compounds

Volatile components analyzed with their respective CAS numbers were purchased from Sigma-Aldrich (Saint Luis, EUA): ethyl acetate (141-78-6), ethyl butanoate (105-54-4), ethyl pentanoate (539-82-2), ethyl hexanoate (123-66-0), ethyl heptanoate (106-30-9), ethyl octanoate (106-32-1), ethyl nonanoate (123-29-5), ethyl ecanoate (110-38-3), ethyl undecanoate (627-90-7), ethyl dodecanoate (106-33-2), diethyl succinate (123,25-1), ethyl lactate (97-64-3), ethyl cinnamate (103-36-6), ethyl anthranilate (87-25-2), ethyl isobutanoate (97-62-1), ethyl 3-hydroxybutanoate (5405-41-4), ethyl isovalerate (108-64-5), ethyl 2-methylbutanoate (7452-79-1), phenylethyl acetate (103-45-70), hexyl acetate (142-92-7), S-furfuryl thioacetate (13678-68-7), furfuryl acetate (623-17-6), isobutyl acetate (110-19-0), isoamyl acetate (123-92-2), 3-methyl-1-butanol (123-51-3), methanol (67-56-1), 1-butanol (71-36-3), 2-butanol (78-92-2), 1-propanol (71-23-8), 2-phenyletanol (60-12-8), 1-hexanol (111-27-3), furfuryl alcohol (98-00-0), propanoic acid (79-09-4), butanoic acid (107-92-6), valeric acid (109-52-4), hexanoic acid (142-62-1), heptanoic acid (111-14-8), octanoic acid (124-07-2), pelargonic acid (112-05-0), decanoic acid (334-48-5), undecanoic acid (112-37-8), 10-undecenoic acid (112-38-9), isobutyric acid (79-31-2), isovaleric acid (503-74-2), α-pinene (7785-70-8), β-pinene (19902-08-0), geraniol (106-24-1), α-terpineol (98-55-5), limonene (5989-27-5), citronelal (2385-77-5), cedrene (469-61-4), γ-nonalactone (104-61-0), β-damascenone (23696-85-7), α-ionone (127-41-3), β-ionone (14901-07-6) and 4-methyl-2-pentanol (108-11-2).

#### Qualitative analysis of SB juice (before-fermentation)

The identification of the fraction of volatile compounds was performed as recently described by Arcari et al. (27). A Varian CP-3900 (USA) gas chromatograph equipped with a Varian Saturn 4000 trap ion mass spectrometer and the Saturn GC-IT / MS version 5.51 Workstation software was used to identify volatile compounds. Chromatographic separation was performed employing a ZB-WAXplus (60 m x 0.25 mm x 0.25 μm) column from Zebron (USA).

### Quantitative analysis of SB fermented (after-fermentation)

Quantification of volatiles was performed as described recently by Arcari et al. (27). Quantitative analyzes were performed on a Thermo Scientific Trace 1310 (USA) gas chromatograph equipped with a flame ionization detector (FID) and ChromQuest software. To determine the contribution of a chemical compound identified in the global aroma of SB fermented the value of olfactory activity (OAV) was determined as an indicator of the importance of a specific compound for the aroma of the sample and was calculated as the ratio of the concentration of the individual compound and the threshold of perception described in the literature (21). Only those components that reached OAV > 1 were considered important for the overall contribution of the aroma and the clustering was represented by the heat mapp as described by Babicki et al. (28).

### Statistical analysis

The assay was carried out in triplicates. With the data collected, it was calculated the mean values, standard deviation, the coefficient of variation, ANOVA/MANOVA, and TUKEY HSD (p≤ 0.05). The effects were considered statistically significant at the p = 0.05 level. Comparisons between strains were calculated using R Core Team (2018). Principal component analysis (PCA) was used to determine the best description and discrimination of the aroma profile between the strains evaluated by R Core Team (2018).

## ACKNOWLEDGEMENTS

The authors are grateful to winery/vineyard owners for allowing us to collect the samples and financial support was provided by Epagri/Ufrgs.

## COMPLIANCE WITH ETHICAL STANDARDS

### Ethical approval

This article does not contain any studies with human participants or animals performed by any of the authors

The sequencing data associated with this study have been deposited in GenBank under accession number: MN998155.1; MN998154.1; MN998153.1; MN998152.1;MN998151.1; MT002913.1; MT002912.1 and (Genbank accessions XX and XX)

