## Supplemental Table1 for "The effect of wild *Saccharomyces* on composition and aroma of the Sauvignon blanc"

### Supplementary information

**TABLE S1**:Volatile compounds identified in grape must Sauvignon Blanc (SB) with their respective retention times, identification ions, retention indices, identification methods, odor descriptor and perception threshold. Nf = not found, STD = mass spectra and retention index in agreement with the standard of the volatile compound. MS = mass spectra in agreement with the spectral database (NIST considering minimum 70% similarity).

| Retention time | Compound | Selected Ions | LTPRI Calculate | LTPRI Literature | Identification Method | Odor Descriptor | Perception Threshold (μg $L^{-1}$) |
|---|---|---|---|---|---|---|---|
| 7,592 | Ethyl acetate | 61, 88 | 891 | 890[e] | STD, MS | Solvent[a],[b], fruity [c,d], balsamic[d] | 12000[e] |
| 10,937 | Ethyl isobutanoate | 116, 88, 71 | 966 | 968[e] | STD, MS | Fruity, banana[h] | 15[e] |
| 17,674 | Ethyl pentanoate | 88, 57, 85 | 1126 | 1132[e] | STD, MS | Fruity, apple[e] | 5[e] |
| 18,193 | 1-butanol | 39, 57, 72 | 1159 | 1165[e] | MS | Medicinal[e] | 150000[e] |
| 18,262 | Thioacetic acid | 43,61, 42 | 1167 | 1163[j] | MS | Toasted[j], onion, garlic[n] | Nf |

| | | | | | | | |
|---|---|---|---|---|---|---|---|
| 19,012 | 3-hexen-2-one | 83, 55, 43 | 1182 | 1211[n] | MS | Boiled vegetables, metal[j] | Nf |
| 21,544 | 2 6-dimethyl-4-heptanone (isovalerone) | 57, 85, 41 | 1195 | Nf | MS | Nf | Nf |
| 24,320 | 3-methyl-1-butanol | 42, 55, 70 | 1202 | 1205[j] | STD, MS | Burnt, alcohol[c,h], nail polish, whiskey[d] | 30000[l] |
| 25,217 | 2-hexanol | 45, 69, 41 | 1217 | 1238[n] | MS | Nf | Nf |
| 34,006 | 1-hexanol | 56, 69, 84 | 1369 | 1372[l] | STD, MS | Herbaceous, greasy[i], resinous; floral, green, cut grass [d, h] | 110[e] |
| 34,592 | (E)-3-hexen-1-ol | 41, 67 | 1376 | 1379[n] | MS | Herbaceous [n] | 70[o] |
| 35,861 | (Z)-3-hexen-1-ol | 67, 41 | 1394 | 1401[n] | MS | Herbaceous, bitter, fatty[e] | 1000[e] |
| 36,653 | 2,4-hexadienal | 81, 39, 41 | 1402 | 1407[n] | MS | Vegetable[n] | 60[o] |
| 37,351 | Isocitronellol | 83, 55, 41 | 1459 | 1488[j] | STD, MS | Candy, roses[j] | 40[o] |

| | | | | | | | |
|---|---|---|---|---|---|---|---|
| 41,250 | Linalool oxide | 59, 43 | 1480 | 1484[n] | MS | Candy, floral, woody[n] | 500[e] |
| 43,139 | Benzaldehyde | 51, 77, 106 | 1493 | 1529[e] | MS | Almonds[e] | 2000[e] |
| 43,587 | Isovaleric acid | 60, 43, 41 | 1666 | 1660[e] | STD, MS | Candy, cheese[k], rancidity[e] | 3000[e] |
| 43,921 | Ethyl decanoate | 88, 101, 29 | 1668 | 1651[e] | STD, MS | Fruity, grape[e] | 200[l] |
| 44,040 | Diethyl succinate | 101, 129, 29 | 1691 | 1690[e] | STD, MS | Wine[c,d,h], toffee[f], fruity[d] | 200000[l] |
| 44,734 | Acetophenone | 105, 77, 51 | 1692 | 1690[n] | MS | Floral, almonds[j] | 65[o] |
| 44,946 | α-terpineol | 81, 136, 43 | 1711 | 1713[e] | STD, MS | Floral, candy[e], anise, mint[j] | 250[l] |
| 44,959 | 1,1,6-trimethyl-1,2-dihydronaphthalene - TDN | 142, 159, 172 | 1697 | 1714[n] | MS | Liqueur[n] | 540[m] |
| 45,552 | 1-decanol | 70, 55, 56 | 1722 | 1735[n] | MS | Candy, fatty[e] | 400[e] |
| 46,065 | Verbenone | 107, 91, 39 | 1725 | 1742[n] | MS | Mint, spices[n] | Nf |

| | | | | | | | |
|---|---|---|---|---|---|---|---|
| 47,829 | 1-undecanol | 55, 69, 41 | 1737 | 1738[e] | MS | Fruity [e], tangerine[j] | 41[e] |
| 61,241 | α-ionone | 136, 121, 93 | 1808 | 1829[n] | STD, MS | Fruity, floral, raspberry, violet [h] | 2,6[l] |
| 61,925 | β-damascenone | 190, 121, 69 | 1815 | 1842[e] | STD, MS | Baked apple[l], floral, honey [d,l] | 0,05[l] |
| 62,388 | Ethyl laurate | 88, 101 | 1838 | 1856[n] | STD, MS | Candy, floral[e], waxy, soap[h] | 1500[e] |
| 63,044 | Hexanoic acid | 60, 73, 41 | 1869 | 1863[e] | STD, MS | Cheese, greasy[e] | 420[l] |
| 63,759 | Decyl isobutyrate | 43, 89, 71 | 1870 | Nf | MS | Nf | Nf |
| 64,386 | Benzyl alcohol | 79, 108, 107 | 1871 | 1874[n] | MS | Candy, fruity[e] | 200000[l] |
| 66,097 | 2-phenylethanol | 65, 91, 92 | 1931 | 1939[n] | STD, MS | Roses, honey[e,k] | 14000[l] |
| 70,677 | Phenol | 94, 66, 65 | 1968 | 1962[n] | MS | Phenolic, medicinal[n] | 5900[o] |

| | | | | | | | |
|---|---|---|---|---|---|---|---|
| 71,382 | β-ionone | 177, 192, 91 | 1985 | 1975[n] | STD, MS | Violeta[d,h,i], balsâmico, rosas[d] | 0,09[l] |
| 72,105 | Isopropyl myristate | 43, 102, 60 | 1999 | 2017[n] | MS | Nf | 800[e] |
| 72,695 | Ethyl myristate | 88, 101, 43 | 2025 | 2044[n] | MS | Lily[j] | Nf |
| 72,881 | γ-nonalactone | 85, 29, 41 | 2032 | 2044[n] | STD, MS | Coconut, peach [b,g,j] | 30[l] |
| 73,384 | Octanoic acid | 60, 73, 43 | 2048 | 2055[n] | STD, MS | Rancidity [d,k], candy, cheese[c], animal, spices[f], unpleasant[d] | 500[l] |
| 80,756 | Ethyl cinnamate | 103, 131, 176 | 2140 | 2139[j] | STD, MS | Honey, cinnamon[c,f], floral, strawberry, plum[f] | 1,1[l] |
| 82,127 | Ethyl palmitate | 88, 101 | 2234 | 2250[n] | MS | Waxy, greasy[e] | 1500[e] |
| 83,100 | Decanoic acid | 60, 129, 172 | 2279 | 2287[e] | STD, MS | Unpleasant[d,k], rancid fat[c], animal[f] | 1000[e] |
| 83,273 | Ethyl-9- hexadecenoate | 55, 88, 69 | 2279 | 2265[n] | MS | Nf | Nf |

| | | | | | | | |
|---|---|---|---|---|---|---|---|
| 87,629 | 2-hexadecanol | 55, 69, 83 | 2310 | 2302[e] | MS | Nf | Nf |
| 96,623 | Hexyl cinnamaldehyde | 129, 117, 91 | 2512 | 2526[n] | MS | Nf | Nf |

Nf = not found, STD = mass spectra and retention index in agreement with the standard of the volatile compound. MS = mass spectra in agreement with the spectral database (NIST considering minimum 70% similarity). [a]Revi et al.(1); [b]Chen; Wang; Xuche (2); [c]Garcia-Carpiteiro et al.(3); [d] Peng et al. (4); [e] Welke et al. (5); [f] Gambetta et al.(6); [g] Pereira et al. (7); [h] Noguerol-Pato et al.(8); [i] Coelho et al.(9);[J] Flavornet , 2016 (10); [m] The Good Scents Company (11); [n]Pherobase, 2016 9 (12); [o]Leffingwell, 2016 (13)

**TABLE S2**: The concentration of fatty acid (µg/L) after fermentation of Sauvignon Blanc.

| Sample | Isovaleric acid | Propanonic acid | Butanoic acid | Pentanoic acid | Hexanoic acid | Isobutyric acid | Heptanoic acid | Octanoic acid | Nonanoic acid | Decanoic acid | Undecanoic acid |
|---|---|---|---|---|---|---|---|---|---|---|---|
| 01PP | 418,07 a | nd | nd | 115,95 ab | 204,49 abc | 2469,61 b | nd | 35294,01 ab | 22,68 c | 313,82 a | nd |
| 06CE | nd | 61,95 a | 22,9 c | nd | 514,82 a | nd | nd | 1939,97 cd | 124,65 ab | 7,26 cd | 34,04 a |
| 11CE | 3,15 b | 18,18 b | 78,25 bc | nd | 383,73 a | nd | nd | 656,85 c | 20,00 c | 2,25 cd | 28,48 a |
| 12M | 423,14 a | 15,27 b | 90,44 bc | nd | 126,68 abcd | 4668,82 b | nd | 41355,92 a | 60,64 cd | 39,26 b | nd |
| 13PP | 689,03 a | 15,87 b | 211,96 a | nd | 225,58 abc | 2709,97 b | nd | 4516,95 cd | 74,55 bc | 38,35 b | nd |
| 26PP | nd | 74,53 a | 141,72 ab | 201,68 a | 93,67 abcd | 1501940 a | 63,07 a | 47158,69 a | 17,23 d | 3,59 cd | nd |
| 28AD | nd | 121,81 a | nd | 14,66 c | 7,24 d | nd | nd | 1144,48 cd | 25,07 c | 0,33 cd | nd |

| | | | | | | | | | | | |
|---|---|---|---|---|---|---|---|---|---|---|---|
| 33CE | nd | 11,25 ab | nd | 15,48 c | 100,40 bcd | nd | nd | 533,61 cd | 18,19 d | 0,30 cd | nd |
| 41PP | 387,82 a | 7,31 c | 91,48 bc | 187,10 a | 351,77 ab | 3554,93 b | nd | 47297,21a | 87,31 bc | 53,92 b | nd |
| SB | 3,15 b | nd | nd | nd | 19,24 cd | nd | nd | 19,60 d | nd | 10,67 c | nd |
| SC2048 | nd | nd | 170,37 ab | 68,03 bc | 121,15 bcd | 2439,60 b | nd | 17624,72 bc | 130,01 a | 51,87 b | nd |
| Mean* | 174,94 | 29,65 | 73,37 | 54,81 a | 195,34 | 2805,66 | 5,73 | 17958,36 | 52,76 | 47,42 | 5,68 |
| p-value | <0,0001 | <0,0001 | 0,0050 | 0,3165 | 0,0013 | 0,0002 | <0,0001 | <0,0001 | <0,0001 | <0,0001 | 0,0003 |

Means followed by the same letter do not differ by Tukey's test considering 0.05 significance level. *obtained considering the nd samples as zero

**TABLE S3**: Concentration of ethyl ester (µg/L) after fermentation of Sauvignon Blanc.

| Sample | Ethyl isobutanoate | Ethyl butanoate | Ethyl 2-methylbutanoate | Ethyl isovalerate | Ethyl pentanoate | Ethyl hexanoate | 3-methyl-1-butanoate | Ethyl heptanoate | Ethyl octanoate | Ethyl decanoate | Diethyl succinate | Ethyl undecanoate | Ethyl laurate | Ethyl lactate | Ethyl cinnamate |
|---|---|---|---|---|---|---|---|---|---|---|---|---|---|---|---|
| SB | 50,24 b | nd | nd | nd | 15,95 a | nd | 2,52 c | nd | nd | nd | 9,95 e | nd | nd | nd | 4,36 b |
| 33CE | 38,11 b | 7,58 d | nd | 51,94 b | nd | 6,87 b | nd | nd | 0,185 d | 0,58 c | 352,81 d | 0,09 c | 6,79 ab | 1138,68 d | nd |
| 28AD | 30,59 b | 7,88 d | nd | 68,07 ab | nd | 5,74 b | nd | 2,58 c | 15,47 b | 0,74 c | 393,49 d | nd | 6,49 ab | 1618,32 d | nd |
| 11CE | 20,28 b | nd | 9,31 b | 49,40 b | nd | 0,11 c | nd | 0,29 c | 0,14 d | 1,83 b | 400,96 d | 0,21 c | nd | 11490,42 b | 4,36 b |
| 06CE | 114,53 b | 14,98 d | nd | 50,97 b | nd | 4,37 c | nd | 0,24 c | 10,00 c | 7,42 a | 1600,93 c | 1,06 bc | 7,58 ab | 7691,16 c | 13,01 a |

| | | | | | | | | | | | | | | | |
|---|---|---|---|---|---|---|---|---|---|---|---|---|---|---|---|
| 13PP | 97,12 ab | 57,44 b | 12,83 b | 464,22 a | nd | nd | 149,69 a | 24,50 b | 214,34 c | nd | 3261,78 bc | 2,00 ab | nd | 4461,40 c | 4,35 b |
| SC2048 | 60,87 b | 23,51 d | nd | 74,49 b | nd | 23,76 c | nd | nd | 526,43 b | 0,46 c | 7186,51 bc | nd | nd | 14727,44 b | nd |
| 12M | 92,07 ab | 53,06 bc | nd | 134,30 b | nd | nd | nd | 193,64 a | 865,29 a | 5,22 ab | 11968,00 b | 3,23 a | 9,30 a | 7368,24 c | 4,36 b |
| 01PP | 62,81 b | 31,69 cd | nd | 65,25 b | nd | 54,14 bc | nd | nd | 299,63 c | nd | 13394,20 b | nd | nd | 3662,88 d | 3,05 b |
| 41PP | 360,28 a | 85,22 a | 91,90 a | 157,16 ab | nd | 117,28 ab | nd | nd | 873,51 a | 2,51 ab | 13859,32 b | 0,60 c | nd | 2382,97 c | 1,43 b |
| 26PP | 46,12 b | 90,10 a | 111,13 a | 96,87 ab | nd | 162,88 a | nd | nd | 1025,26 a | nd | 26709,35 a | nd | nd | 39063,35 a | 4,57 b |
| Mean | 88,45 | 33,77 | 20,47 | 110,24 | 1,48 | 34,10 | 13,84b | 20,11 | 348,20 | 1,71 | 7194,30 | 0,65 c | 2,74 | 8509,53 | 3,59 |
| p-value | 0.0036 | <0.0001 | <0.0001 | 0.0258 | <0.0001 | <0.0001 | 0.0027 | <0.0001 | <0.0001 | 0.0025 | 0.0001 | 0.0041 | 0.0309 | 0.1668 | 0.6426 |

Means followed by the same letter do not differ by Tukey's test considering 0.05 significance level. *obtained considering the nd samples as zero

10. Flavornet and human odor space. ⟨http://www.flavornet.org/flavornet.html⟩ (accessed on 23.03.16).

11. The Good Scents Company Information System. ⟨http://www. thegoodscentscompany.com⟩ (accessed on 23.03.16).

12. Pherobase Database of Pheromones and Semiochemicals. http://http://www.pherobase.com/ (accessed on 23.03.16).
